# Sloppiness and Action Constraint in Cell State Transitions: Are Single Cells Sloppy?

**DOI:** 10.64898/2025.12.31.697145

**Authors:** Yuxuan Wang, Junda Ying, He Xiao, Miao Huang, Lei Zhang, Weikang Wang

## Abstract

Single-cell dynamics during cell state transitions (CST) are highly constrained, enabling precise control. However, there are challenges in achieving a system-level understanding or extracting general principles for CST dynamics due to the complexity of gene-gene interactions. Here, we introduce a new perspective to deal with these challenges using Fisher information. We found that, during CST, single cells exhibit pronounced sloppiness: cell states are sensitive only to a few “stiff” parameters while remaining robust to changes in numerous “sloppy” parameters. Critical transitions coincided with changes in stiff parameters. Moreover, stiff parameters typically exhibited minimal fluctuations and low velocities, whereas sloppy parameters allowed greater flexibility. Together, these findings can be summarized by stating that transition paths approximately adhere to a principle of least action. By characterizing the low dimensionality and constraints of CST through sloppiness and action, our work thus introduces a new conceptual and computational framework for analyzing single-cell dynamics.

**Teaser:** Except when transitions occur, single-cell dynamics exhibit sloppiness, adhering approximately to a least action principle.

## Introduction

Understanding the dynamics of cell state transitions (CST), a phenomenon prevalent in processes like development and metastasis, is pivotal in biology. High-throughput methods such as single-cell RNA sequencing (scRNA-seq) have significantly expanded our knowledge of this complex process (*1-3*). However, the paradox of CST dynamics lies in how cells achieve remarkably precise and reproducible outcomes despite the complexity and noise inherent in underlying molecular processes; a system-level understanding, particularly the physical principles guiding CST dynamics, remains elusive (*4*).

Researchers often interpret this paradox through the concept of a latent manifold, a low-dimensional surface emerging from underlying gene regulatory networks (GRNs) that acts as a physical constraint, effectively “funneling” cells into a limited set of cell fates (*4-7*). The Waddington landscape metaphor is one of the most renowned illustrations that aligns with this concept. Various methods have been developed to quantify this landscape using scRNA-seq data (*8-11*). The inference of RNA velocity sparked studies to derive governing equations for exploring the driving forces of CST dynamics (*12, 13*). Additionally, the principle of least action in classical dynamics has led to speculation about its applicability in calculating transition paths in CST dynamics (*14-16*). However, these studies usually rely on low-dimensional embeddings such as UMAP or diffusion maps (*17, 18*), which can distort local distance metrics when projected into two or three dimensions (*19*). Furthermore, none of the previous studies have explicitly defined the constraints imposed by the latent manifold.

Unlike these methods, Fisher information, along with information geometry, offers an alternative lens for characterizing the low-dimensional emergent behaviors (*20, 21*). For instance, Fisher information has already been applied to the study of neural manifolds (*22-24*).

The application of Fisher information lies in its dual functions: serving as a Riemannian metric and a measure of parameter sensitivity. First, as a specific type of Riemannian metric, Fisher information can be used to explore the geometric properties of manifolds. On Riemannian manifolds equipped with the Fisher information metric, the action of transition paths can be readily calculated. Specifically, the integrand in the action integral, known as information velocity—defined as the product of the parameter velocity (time derivative) and the Fisher information—measures the variation rate of the probability distribution. These concepts have been applied to study thermodynamic length and information change in non-equilibrium processes (*25-27*). And the information velocity analysis can be extended to study CST dynamics using our previously developed framework SCIM (single cell information manifold) (*20*).

Second, by quantifying the sensitivity of probability density functions to control parameters, Fisher information allows us to define sloppiness. Sloppiness describes the phenomenon in multi-parameter models where only a few parameter combinations significantly impact the model’s fit to collective behaviors, providing an alternative understanding of robustness (*21, 28, 29*). And the model is particularly sensitive to those influential combinations, known as stiff parameters, while the rest are considered sloppy parameters (*21, 29*).

In a multi-parameter system, sloppiness is usually assessed by examining the eigenvalue spectrum of the Fisher information matrix (FIM). The eigenvalues of the FIM quantify the impact of changes in parameter combinations along their corresponding eigenvectors. The eigenvectors corresponding to stiff parameters are constrained by the observed data. This results in a model manifold of predictions, with coordinates as model parameters, constrained within a low-dimensional hyper-ribbon structure (*21, 29-32*). For example, Gutenkunst et al. identified that eigenvalue spectrum of the FIM in systems biology models exhibit such a structure (*33*). Stiff and sloppy parameters may be associated with the plasticity and robustness of cell states and functions, respectively (*28*).

Beyond characterizing the “static” parameter structure, sloppiness can be extended to the analysis of neural dynamics and constraint on circuit parameters (*34*). For instance, Panas et al. fitted the network activity of cultured rat hippocampal neurons using pairwise maximum entropy model and discovered that the model is sloppy. They also found that neurons with sloppy parameters exhibited faster and larger fluctuations compared to those associated with stiff parameters (*35*). Similarly, Adrian Ponce-Alvarez et al. investigated the neural population activity in the auditory cortex of anesthetized rats and found that stimulus-evoked activity predominantly occurs along sloppy dimensions, while cortical state transitions progress along stiff directions (*36*).

Viewing CST as an adaptation process, the cell system adjusts its parameters to improve fitness, akin to an online model fitting to a changing environment (*37*). The parameters of this model may be attributed to the regulation of different genes, in response to external signals across various stages during CST (*38-40*). Identifying stiff and sloppy properties of these parameters at different stages offer an alternative approach to understand the regulatory mechanism underlying CST. However, few studies have explored single-cell dynamics from this perspective. In this work, we examined the sloppiness in this dynamic fitting processes of cellular systems and developed a framework for analyzing CST dynamics in parameter space. Our findings reveal that cellular systems exhibit significant sloppiness, with stiff parameters varying as cells approach transition states. Notably, the transition paths of CSTs on information manifolds roughly follow a principle of least action.

## Results

### Framework for analyzing sloppiness of CST with FIM

In this study, we fulfilled fitting a special probabilistic model from the scRNA-seq data and constructed a corresponding statistical manifold for CST using SCIM (Methods). Each cell is embedded as a multi-variate Gaussian distribution (Fig. 1A). The distribution parameters serve as coordinates on this manifold, while the FIM defines the Riemannian metric (Fig. 1B). Intuitively, the embedded distributions capture the uncertainty or neighborhood diversity of cells. First, we can analyze the eigenvalue spectrum of the FIMs across different cells with this fitted model (Fig. 1C). The spectral analysis uncovers the sloppiness of individual cells. Large eigenvalues indicate stiff parameters, suggesting critical regulatory mechanisms. And the distribution of eigenvalues and eigenvectors in cell cluster reveals the heterogeneity among single cells.

**Fig. 1.**
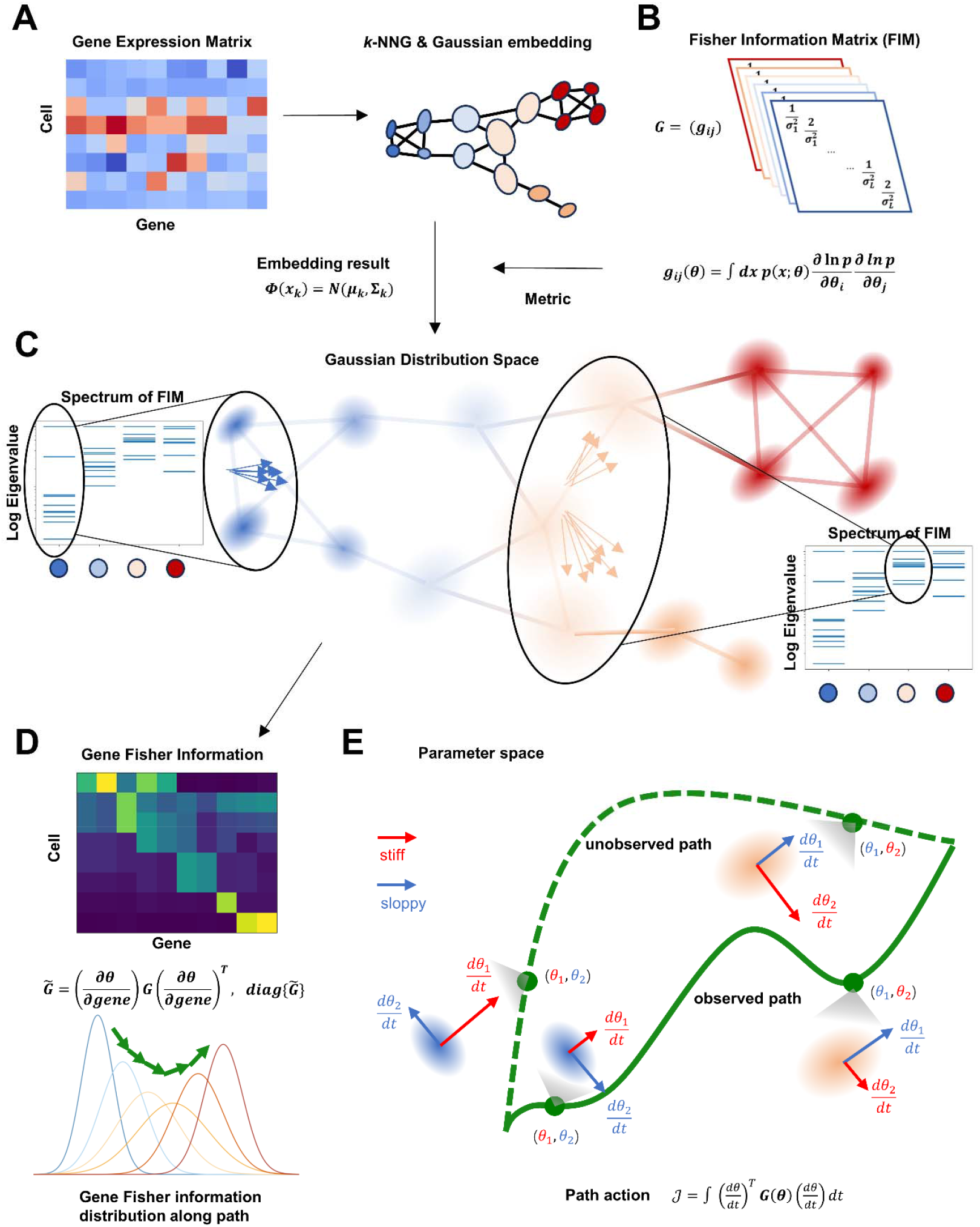
Analyzing single cell state transition with spectrum of FIM. (**A**) Through building the cell *k*-nearest neighbor graph (*k*-NNG) and Gaussian embedding, SCIM maps each cell in scRNA-seq data into a multivariate Gaussian distribution. (**B**) Each single cell is assigned with a FIM which is regarded as the Riemannian metric of the cell state transition manifold. (**C**) Each shadowed ellipse denotes the embedded Gaussian distribution of single cell. The FIM eigenvalue spectrum of different cell clusters are displayed in the box. The arrow bundles in the black circles represent the eigenvectors corresponding to the largest eigenvalue for all cells within each cluster. (**D**) The distribution of genes’ Fisher information is used to assess the potential regulatory mechanisms in CSTs. The curves represent the variation of gene Fisher information distribution along the transition path (color indicates the stage of transition). (**E**) Action constraint on corresponding the parameter Fisher information and the parameter velocity along the transition paths is illustrated. Each point represents a Gaussian distribution corresponding to the parameters, with local ellipses indicating stiff (red) and sloppy (blue) directions in parameter space. Arrow length signifies the magnitude of parameter velocity. Compared with the hypothetical or unobserved path (dashed green curve), the observed path (solid green curve) is more constrained along the stiff directions.

Second, besides analyzing the static spectrum, we can analyze the variation in the FIM eigenvalue spectrum along the estimated transition path (Methods). The changes in the spectrum along the path reveal variations in the parameter space beyond dynamics space. Specifically, an increase in stiff parameters indicates that a cell becomes sensitive to more factors (Fig. 1C). And these factors can be analyzed by examining the corresponding eigenvectors. These results reflected how the key parameters controlling the cell state dynamically change during CST and may associate with cell plasticity.

Third, we can calculate the Fisher information for each gene through coordinate transformation (Methods) and investigate the variation in genes’ sloppiness and stiffness during CST (Fig. 1D). The distribution of genes’ Fisher information changes along the transition path, enabling the analysis of key markers and regulatory genes at different transition stages.

Finally, we can discern common characteristics and explore the general principles guiding CST dynamics through analyzing transition paths on the obtained manifolds across different datasets. Even though these “observed” transition paths are inferred, they still capture underlying real-world dynamics, compared to those that remain hypothetical or unobserved. Specifically, we calculated parameter velocity using RNA velocity and further examined the corresponding information velocity and action on the manifold along these transition paths on the manifolds. The variations in parameter velocities of stiff and sloppy parameters not only reflect critical transitions but also reveal action constraints on CST dynamics (Fig. 1E).

### Analysis of the FIM eigenvalue spectrum reveals sloppiness of CST

We applied our method on several scRNA-seq datasets including zebrafish pigmentation (*41*) and dentate gyrus neurogenesis (*42*). During zebrafish pigmentation, proliferating progenitors differentiate into pigment progenitors and Schwann cell precursors, ultimately forming melanophores, xanthophores, and Schwann cells, representing a tri-fate differentiation process (Fig. 2A, left). The steady states consist of the initial phenotype (proliferating progenitors) and the terminal phenotypes (melanophores, xanthophores, and Schwann cells), while the transient states include pigment progenitors and Schwann cell precursors. Pigment progenitors act as branch points, differentiating into multiple pigment cell types, whereas Schwann cell precursors develop exclusively into Schwann cells, representing a single-lineage critical transition. Dentate gyrus neurogenesis encompasses the differentiation of multiple cell types. For comparison with multi-lineage CSTs like zebrafish pigmentation, we focused on the transition from radial glia to granule cells via neuroblasts, a single-lineage CST process (Fig. 2A, right). The critical transition occurs at the transient state, i.e., neuroblasts.

**Fig. 2.**
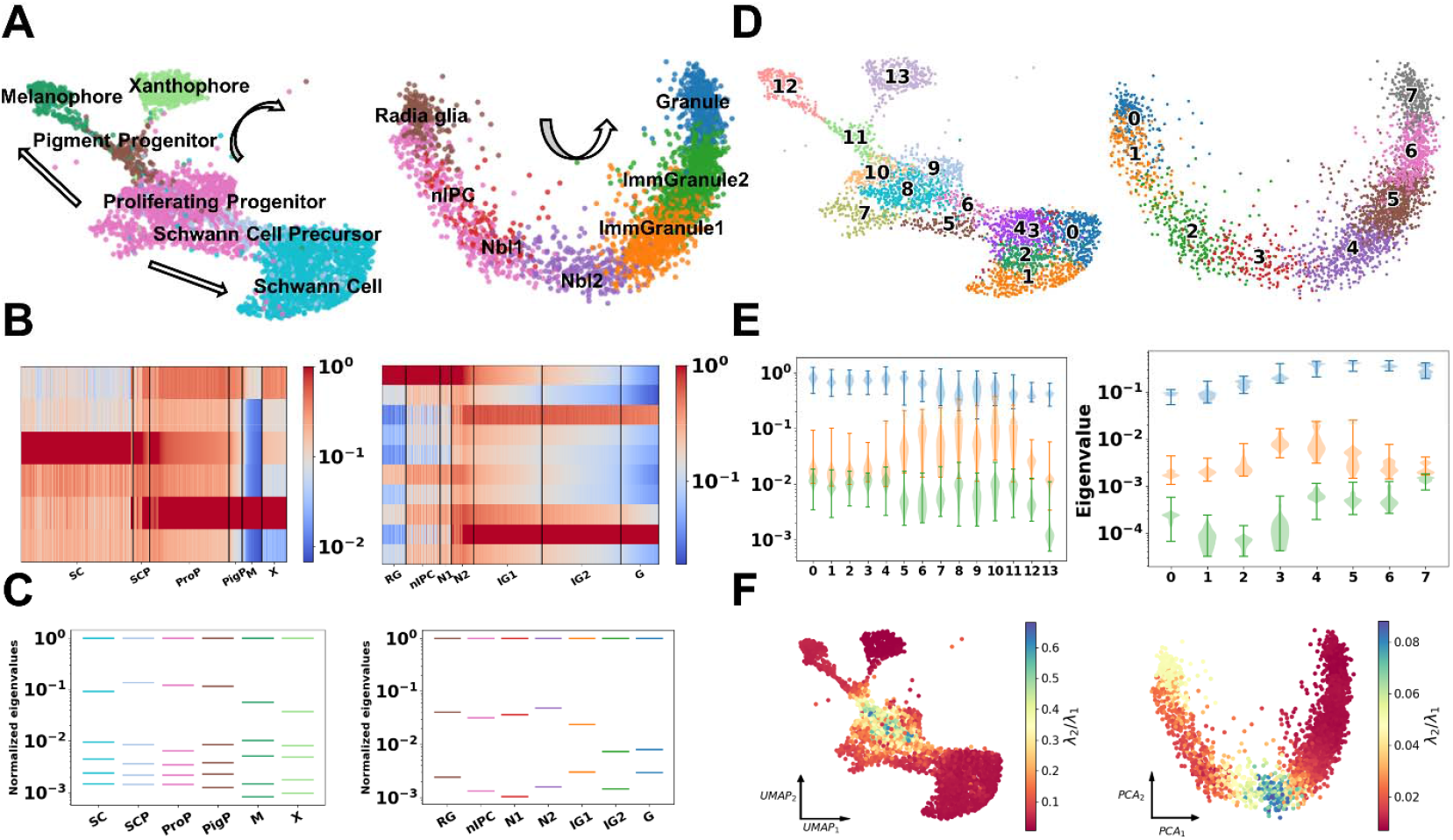
Spectrum analysis of the Fisher information matrix. (**A**) Cell type annotation of zebrafish pigmentation dataset (left) and dentate gyrus neurogenesis dataset (right). Arrow represents the direction of CST. (**B**) FIM spectrum of (***μ, σ***) for zebrafish pigmentation (left) and dentate gyrus neurogenesis (right) (only ***σ*** are shown for simplicity). Each row represents a latent parameter, while each column represented a cell. The color represented the normalized Fisher information of the parameter in that cell. SC, Schwann cell; SCP, Schwann cell precursor; ProP, Proliferating progenitor; PigP, Pigmentation progenitor; M, Melanophore; X, Xanthophore; RG, Radial Glia; nIPC, nIPC; N1, Nbl; N2, Nbl2; IG1, ImmGranule1; IG2, ImmGranule2; G, Granule. (**C**) Averaged eigenvalue spectrum of FIM for each cell type in zebrafish pigmentation (left) and dentate gyrus neurogenesis (right). The bars represent the eigenvalues averaged across individual cells normalized by the largest eigenvalue. The colors of the bars match the dot color of cell type in panel (A). (**D**) PAGA clustering of zebrafish pigmentation (left) and dentate gyrus neurogenesis (right). Colors represent different clusters. (**E**) Violin plot of eigenvalue distribution of Hotspot FIM in each cluster for zebrafish pigmentation (left) and dentate gyrus neurogenesis (right). In each cluster, blue, orange and green violins represent the distribution of the 1^st^, 2^nd^, 3^rd^ eigenvalues, respectively. These values were not normalized. (**F**) The ratio of the 2^nd^ to the 1^st^ eigenvalues of the Hotspot FIM for zebrafish pigmentation (left) and dentate gyrus neurogenesis (right).

With SCIM, we obtained an *L*-dimensional embedding represented by the parameters (means and variances; *μ*and *σ*) of a multivariate diagonal Gaussian distribution for each cell. The FIM of Gaussian distribution with diagonal covariances can be written analytically. For 1-D Gaussian distribution, the Fisher information of *μ* and *σ*are 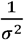 and 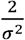 respectively (*43, 44*). The FIM (denoted as *FIM*_*μ,σ*_) of a *L*-dimensional diagonal Gaussian distribution is

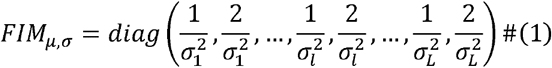

We first investigate the FIM spectrum of each single cell within (***μ***,***σ***)coordinate. Since the Fisher information of *μ* is one half of that of corresponding *σ*, we can focus on analyzing either ***μ*** or ***σ*** in practice. Our findings indicate that Fisher information can effectively characterize cell types (Fig. 2B). For example, proliferating progenitors and Schwann cells exhibit significantly different Fisher information values, with notable differences in the dimensionality with the largest Fisher information (Fig. 2B left). In neurogenesis, the neuroblasts (neuroblast2) display apparent signature of transition. As transition proceeds, cell types like radial glia and neuronal intermediate progenitor (nIPC) exhibit largest Fisher information in the first dimensionality of (***μ***,***σ*)** coordinate (first row in Fig. 2B right) and cell types like immature granule cells and granule cells have largest Fisher information in the ninth dimensionality of (***μ***,***σ*)** coordinate (ninth row in Fig. 2B right). But in neuroblast2, some cells have the largest Fisher information in the first dimensionality, while some cells have the largest Fisher information in the ninth dimensionality, indicating the transitory role of neuroblast2 (Fig. 2B right). This characterization is consistent with the geometric analyses demonstrating that low coarse Ricci curvature (CRC) regions delineate transition states (Fig. S1)(*20*).

However, the analysis in the (***μ***,***σ*)** coordinate has several drawbacks. One issue is that Gaussian embedding is derived from training a neural network, which introduces randomness. Consequently, interpreting sloppiness using the Fisher information of (***μ***,***σ*)** depends on the embedding results (*20*). Another is that *μ* and *σ* lack biological interpretability. Fortunately, the geometry properties of the manifold obtained from SCIM, such as curvature, remain invariant (*20*). This invariance allows us to treat different (***μ***,***σ*)** as coordinate transformations, representing various coordinate systems while preserving the manifold geometry. To overcome the issue mentioned above, we performed the coordinate transformation to convert (***μ***,***σ*)** parameters into gene module parameters, such as Eigengenes and Hotspot (*45, 46*). Eigengene represents the first principal component (PC1) of gene modules identified through gene correlation clustering, while Hotspot modules are generated by identifying genes that share informative co-variation and similarity in proximal cells (Methods). We performed statistics on the eigenvalue spectrum of Eigengene and Hotspot FIM. On a logarithmic scale, both the mean (Fig. 2C) and distribution (Fig. S2A, B) of the Eigengene FIM eigenvalues exhibit sparse patterns and substantial differences in magnitude for each cell type across different datasets, indicating only a few parameter combinations are stiff, while the majority are sloppy. The spectrums of the Hotspot FIM exhibit similar patterns (Fig. S2C, D). This consistency further supports the conclusion that individual cells exhibit sloppiness during CST.

These findings align with the widely accepted notion of low intrinsic dimensionality and are consistent with the eigenvalue spectrum of the FIM in complex model fitting (*21, 29, 33*). Notably, the number of stiff parameters is less than the SCIM embedding dimension *L* and remains robust by changes in this hyperparameter (Fig. S3, Fig. S4).

### Increase of stiff parameter at transitory stages

We further divided the entire cell population into clusters at a higher resolution using PAGA (Fig. 2D) and examined the dominant parameter combinations in each cluster (*41, 47*). The system is primarily influenced by stiff parameters, as defined by the eigenvectors of the FIM corresponding to similarly large eigenvalues. An increase in the number of stiff parameters suggests a higher degree of freedom and greater cell plasticity. The corresponding eigenvectors may associate with various regulatory or functional modules given that Eigengene and Hotspot modules both reflect properties of GRN.

We found that across both zebrafish pigmentation with multiple lineages and dentate gyrus neurogenesis with a single lineage, the gap between the first and second eigenvalue significantly narrows at transitory stages (Fig. 2E). In single-lineage dataset, increased stiffness of more parameter combinations near transitory stage indicates enhanced gene interactions and dominant gene module switching during CST. In multi-lineage dataset, the gap between the first two eigenvalues becomes smaller in undifferentiated clusters, occasionally resulting in overlaps in the corresponding distributions (cluster 8-11). This suggests that dominant gene module selection occurs during cell fate decisions, with only certain modules becoming dominant during specific differentiation processes once cell fate is partially established. These results are robust, with similar phenomena observed in the FIMs of Eigengene and Hotspot under various hyperparameter settings (Fig. S3, Fig. S4).

We also examined the ratio between the second and first eigenvalue (λ_2_/λ_1_) for each cell under Hotspot coordinates (Fig. 2F). Consistent with the violin plot results, these ratios peak in the same regions. This suggests that biological systems exhibit sloppiness at single-cell resolution, with the switch and selection of stiff parameter combinations occurring on a single-cell level. These findings remain consistent in the FIMs of Eigengene and Hotspot across various hyperparameter settings (Fig. S5, Fig. S6). The analysis of pancreatic endocrinogenesis dataset can be found in Fig. S7 (*48*).

### Variation in genes’ sloppiness during CST

The sloppiness analysis of the CST system can be further extended to the gene level. Through coordinate transformation, we obtained the gene-by-gene FIMs. The Fisher information for individual genes are diagonal elements in these FIMs. As demonstrated in Fig. 3A, the average values and variances of normalized Fisher information for selected genes vary across different cell types in zebrafish pigmentation, reflecting the variation in gene stiffness among different cell types.

**Fig. 3.**
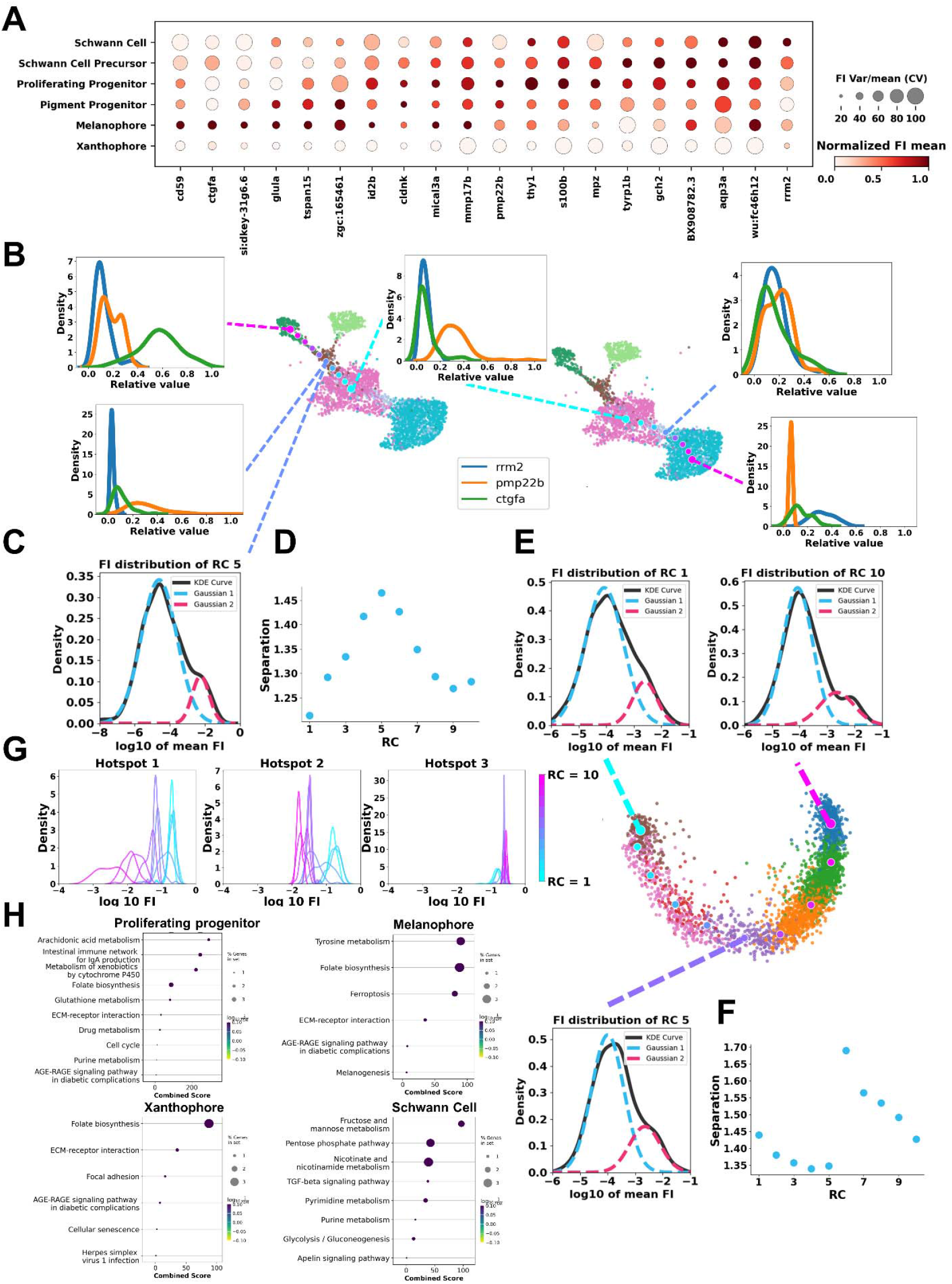
The Gene Fisher information analysis of CST processes. (**A**) Dot plot of the Fisher information of the 20 selected genes grouped by cell types along simulated transition paths. The dot size represents the coefficient of variation (CV) of the gene Fisher information in the cell type, and the color represents the mean gene Fisher information in the cell type, which is normalized gene-wise. The selected genes are the 20 genes with the largest mean Fisher information. (**B**) The variation of individual gene Fisher information along RCs. Two transition paths (left: Proliferating Progenitor to Melanophore; right: Proliferating Progenitor to Schwann Cell) were plotted (Method). The Fisher information of each gene was normalized by the maximal Fisher information of that gene across all cells. Small dots represent single cells with color indicating cell type. Large dots represent RCs with color denoting the direction of transition (from cyan to magenta). (**C**) The distribution of average Fisher information for all genes at the 5^th^ RC from proliferating progenitor to melanophore. (**D**) Variation in the separation of two peaks in the distribution along the transition paths from proliferating progenitor to melanophore. (**E**) Variation of individual gene Fisher information distribution during the transition in the dentate gyrus neurogenesis dataset. Small dots represent single cells with color indicating cell type. Large dots represent RCs with color denoting the direction of transition (from cyan to magenta). (**F**) Variation in the separation of two peaks in the gene Fisher distribution during the transition in dentate gyrus neurogenesis. (**G**) Shift in the Hotspot Fisher information distribution along the transition from proliferating progenitor to melanophore. (**H**) The Gene enrichment analysis of the top 20 genes with the largest gene Fisher information in specific cell type (proliferating progenitor, melanophore, xanthophore, and Schwann cell).

Utilizing RNA velocity, we calculated the average transition paths in all the analyzed datasets (Methods) (*40*). The average transition path is discretized into multiple reaction coordinates (RCs) (Fig. 3B, center). RCs are a series of points evenly distributed along the transition path, and the multi-dimensional state space is divided into Voronoi grids (*40, 49*). In the context of CST, RC is conceptually similar to pseudo-time trajectory in single-cell genomics (*1, 3*).

We examined how the distribution of individual gene’s Fisher information varies, normalized by each gene’s maximum value across the entire population, for cells residing within the same Voronoi grid (i.e., sharing the same RC index) along transition paths (paths from proliferating progenitors to melanophores and Schwann cells). For instance, the Fisher information distribution of *rrm2* becomes more dispersed, indicating that *rrm2* becomes stiffer in some cells when transiting into the Schwann cell. The results of the transition path from proliferating progenitors to xanthophores are illustrated in Fig. S8.

We also calculated the average Fisher information (log scale) for all genes within the corresponding Voronoi grid of each RC along the transition paths. We found that the variations in this distribution exhibit similar dynamic patterns. As shown in Fig. 3C, the distributions at most RCs are approximately bimodal, suggesting that a small proportion of genes have high Fisher information. We used a two-component Gaussian mixture model to fit each distribution (Fig. 3C) and found that the separation between the two components peaks during CST (Fig. 3D). These peaks are located around transitory cell states such as pigment progenitors and Schwann cell precursors in the zebrafish pigmentation, suggesting that the genes with high Fisher information become relatively stiffer during the transitory stage. Similar result is observed at the transitory stage in the single-lineage dentate gyrus neurogenesis dataset (Fig. 3E, F). In conjunction with the results of separation, the peaks in the Gini coefficient also reflect a “stiff get stiffer” phenomenon in Fisher information, suggesting changes in overall gene sloppiness during the transitory stage (Fig. S9).

Similar to that of individual genes, we also observed the shift of distribution of individual Hotspot module’s Fisher information along the transition path (Fig. 3G). We then conducted a gene ontology (GO) enrichment analysis on the genes whose sloppiness varies significantly in the zebrafish dataset (Fig. 3H). This analysis revealed that these genes are related to several critical biological pathways associated with pigmentation and Schwann cell development, including tyrosine metabolism, Notch signaling, and melanogenesis.

### Increase of stiff direction at transitory stages

The variation in the FIM eigenvalues reveals that more parameters become stiff as cells approach the transitory stage. Building on this, we further examined the variation in FIM eigenvectors, particularly the one corresponding to the largest eigenvalue. This vector represents the stiffest parameter combination, which we refer to as stiff direction. By analyzing the stiff directions under different biologically meaningful module coordinates and within various cell types, we can glean insights into how CSTs are regulated by underlying GRNs.

We analyzed the pairwise correlation distance between the stiff direction of the Eigengene FIM across different cells (Fig. 4A). Since the stiff direction is a multi-dimensional vector that can have two opposite orientations, the orientation can significantly affect the distance value. We aligned the directions by solving an integer programming problem using the Gurobi optimizer (Fig. S10) and performed clustering on the aligned stiff directions based on correlation distance (Methods) (*50*). For example, in pancreatic endocrionogensis dataset, Fev+ cells were clustered into three groups (Fev+_1, Fev+_2, Fev+_3), but Ngn3-low progenitor cells were assigned to a single group (Fig. 4A, inset heatmap). For better visualization, we mapped the stiff directions to low-dimensional embeddings such as UMAP (Methods) and plotted the vectors of stiff directions for each cell type (Fig. 4A, inset arrow bundles). The distribution of stiff directions is notably narrow at the initial and terminal stages but becomes more dispersed in the transitory region.

**Fig. 4.**
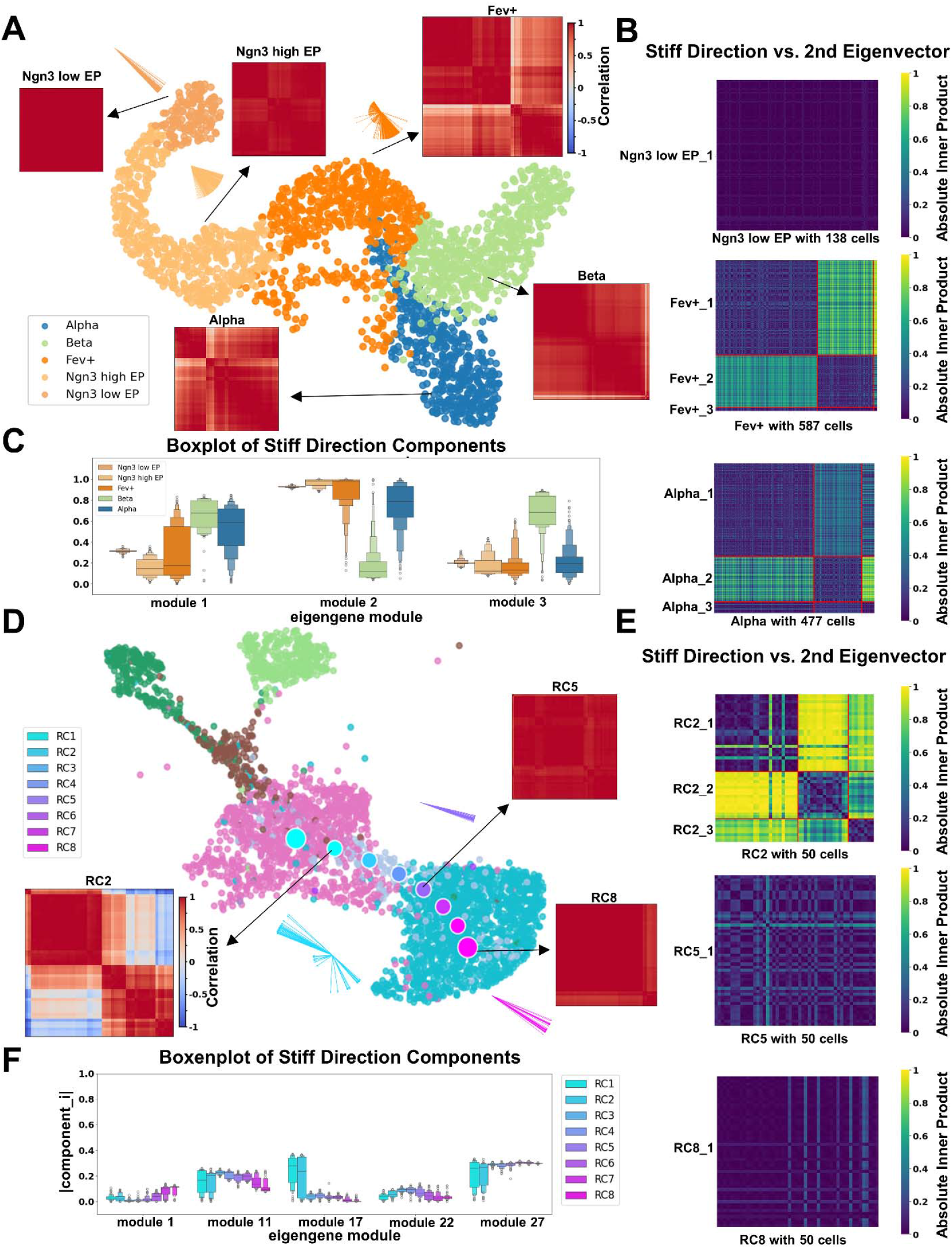
Analysis of stiff directions in CST. (**A**) Distributions of the stiff directions of different cell types in pancreatic endocrinogenesis dataset. The matrix heatmaps display the pairwise correlation distance matrices of the stiff directions within the corresponding cell types. The inserted arrow bundles visually represent the stiff directions (Methods). The arrow lengths are proportional to the size of cell types. Small dots represent single cells with color indicating cell type. (**B**) Similarity between the second eigenvectors and the stiff directions for different cell types. The red line represents the divisions based on the clustering results in panel (A). (**C**)Boxen plots of the absolute magnitudes of the Eigengene component in the stiff directions in different cell types in endocrinogenesis. Successive boxes represent increasing quantiles (percentiles) of the distribution, providing a detailed view of the data density and tails compared to standard box plots. The central line represents the median. (**D**) Same as that in panel (A) except for different RCs along the transition path from proliferating progenitor to Schwann cell in zebrafish pigmentation. Small dots represent single cells with color indicating cell type. Large dots represent RCs with color denoting the direction of transition (from cyan to magenta). (**E**) Same as that in panel (B) except for different RCs along the transition path from proliferating progenitor to Schwann cell in zebrafish pigmentation. (**F**) Same as that in panel (C) except for different RCs along the transition path from proliferating progenitor to Schwann cell in zebrafish pigmentation.

We previously observed a relative increase in the second largest eigenvalue of the FIM at the transitory stage (Fig.2E, F and Fig. S7). Here, we examined the similarity between the second eigenvector and the stiff direction by computing their inner product within each cell type (Fig. 4B). In Ngn3-low progenitor cells, the overall inner product values are low (Fig. 4B top). However, in Fev+ and glucagon-producing α-cells, there are non-diagonal blocks with values close to 1. Notably, the stiff directions and second eigenvectors in Fev+_1 cells correspond approximately to the second eigenvectors and the stiff directions in Fev+_2 cells, respectively (Fig. 4B, middle and bottom). This indicates that the stiff directions in Fev+ cells switch between two types of vectors. Such an observation suggests the occurrence of eigenvalue crossing, where the stiff direction changes abruptly due to a shift in the order of the eigenvalues. Biologically, this corresponds to an increase in the number of dominant directions governing the biological process at this transitory stage.

The elements of the stiff directions in the Eigengene FIM represent the weights of the corresponding eigengene modules. We compared the compositions of Eigengene (absolute weight) in stiff directions (Fig. 4C) (*51*). At the transitory stage (Fev+ cell), the distribution is relatively dispersed across the components, whereas it is more concentrated at the initial and terminal states. Different branches exhibit distinct component distributions; for instance, glucagon-producing α-cells are higher in module3 and lower in module1, while glucagon-producing β-cells show the opposite pattern. The variation in component distribution reflects changes of dominant modules during CST. We conducted the same analyses using Hotspot and obtained similar results (Fig. S11).

We examined the stiff directions along the transition path, discretized by RC (Fig. 4D). In the transition from proliferating progenitors to Schwann cells in zebrafish pigmentation, the correlation distance values between the first stiff directions at RC2 span a large range, whereas the distance values at RC5 and RC8 are nearly equal to 1 (Fig. 4D, inset heatmap). The vector bundles of the first stiff direction on UMAP at RC2 are more dispersed than those at RC5 and RC8. The inner products of the second eigenvectors and the stiff directions reveal the fluctuation in the order of the eigenvalues at RC2 (Fig. 4E), reflecting the heterogeneity at RC2.

By analyzing the compositions of Eigengene in the stiff direction along the RCs (Fig. 4F), we found that several Eigengene modules show dispersive distributions at RC1 and RC2, but maintain either low value (module 17) or high value (module 27) at the following RCs. In contrast to pancreatic endocrinogenesis, where the stiff directions become dispersive at the cell type like Fev+ at the transitory stage (Fig. 4A, B), the dispersive stiff directions at RC1 and RC2 (proliferating genitors) may indicate that the timing of cell-fate decision occurs earlier in zebrafish pigmentation. Analyses of the transitions to melanophores and xanthophores are provided in Fig. S12 and Fig. S13.

### Fluctuations of stiff and sloppy parameters along transition path

Within the SCIM framework, each individual cell is assigned a unique set of parameters. CSTs involve not only changes in the Fisher information of these parameters but also variations in these parameters themselves. In this section, we analyzed the fluctuations of the parameters (eigenvectors of FIM in PCA coordinate) along the transition paths in different datasets (Fig. 5A) using the squared coefficient of variation (*CV*^2^; see Methods).

**Fig. 5.**
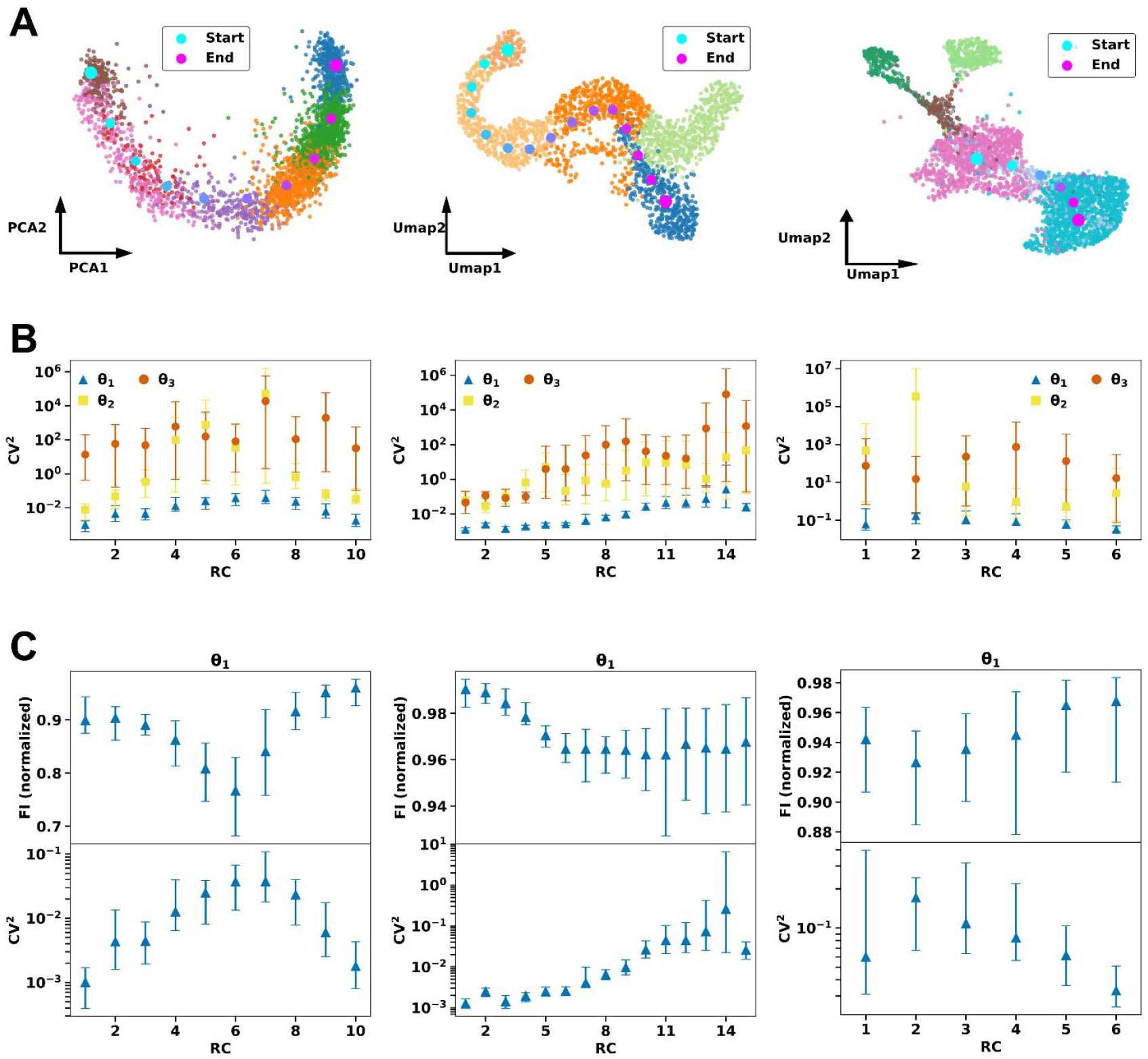
Fluctuation of stiff and sloppy parameters in CST. (**A**) Transition paths discretized by RC for three datasets. Left: dentate gyrus neurogenesis dataset. Middle: glucagon-producing α-cell branch of the pancreatic endocrinogenesis dataset. Right: melanophores branch of the zebrafish pigmentation dataset. Small dots represent single cells with color indicating cell type. Large dots represent RCs with color denoting the direction of transition (from cyan to magenta). (**B**) Squared coefficient of variation () for the parameters corresponding to the 1^st^ (blue), 2^nd^ (yellow), 3^rd^ (orange) largest eigenvalues of the PCA FIM at each RC poin along the transition paths. Left: dentate gyrus neurogenesis dataset. Middle: glucagon-producing α-cell branch of the pancreatic endocrinogenesis dataset. Right: melanophores branch of the zebrafish pigmentation dataset. (**C**) Along the transition path, Fisher information of the stiffest parameter exhibits negative correlation with their. The Fisher information is sum-normalized across all parameters within a single cell. Left: dentate gyrus neurogenesis dataset. Middle: glucagon-producing α-cell branch of the pancreatic endocrinogenesis dataset. Right: melanophores branch of the zebrafish pigmentation dataset.

First, we observed that the stiffest parameters exhibit minimal fluctuations across various transition processes, including dentate gyrus neurogenesis (Fig. 5B, left), pancreatic endocrinogenesis (Fig. 5B, middle), and zebrafish pigmentation (Fig. 5B, right). Second, along each transition path, there is a pronounced negative correlation between the *CV*^2^ and the Fisher information of the stiffest parameters (Fig. 5C). And these observations are robust across different hyperparameter settings (Fig. S14, Fig. S15) and are consistently reproduced in other branches of the pancreatic endocrinogenesis and zebrafish pigmentation datasets (Fig. S16).

Considering that Fisher information characterizes the sensitivity of a parameter, these results imply that the magnitude of fluctuations decreases as parameter sensitivity increases. Similar to multi-parameter models and neural dynamics, low-level fluctuations reflect the constraint imposed on stiff parameters (*21, 29, 34, 36*). Biologically, such parameters may correspond to key genes or critical regulatory mechanisms governing CSTs. Small perturbations in these parameters can lead to significant changes in cell state, prompting the system to regulate them carefully, which may account for the reduced fluctuations observed in our analyses.

### Least action constraint in CST

Following analyses on the parameter fluctuations, we further examined the velocities (time derivative) of stiff and sloppy parameters in each single cell. We calculated the parameter velocity of (***μ***,***σ*)** based on RNA velocity (Methods) and found a clear negative correlation between Fisher information and parameter velocity (Fig. 6A-C). For instance, in dentate gyrus neurogenesis, the Fisher information of σ_1_ is low at the initial stage and high at the final stage, whereas its parameter velocity shows an opposite trend (Fig. 6A).

**Fig. 6.**
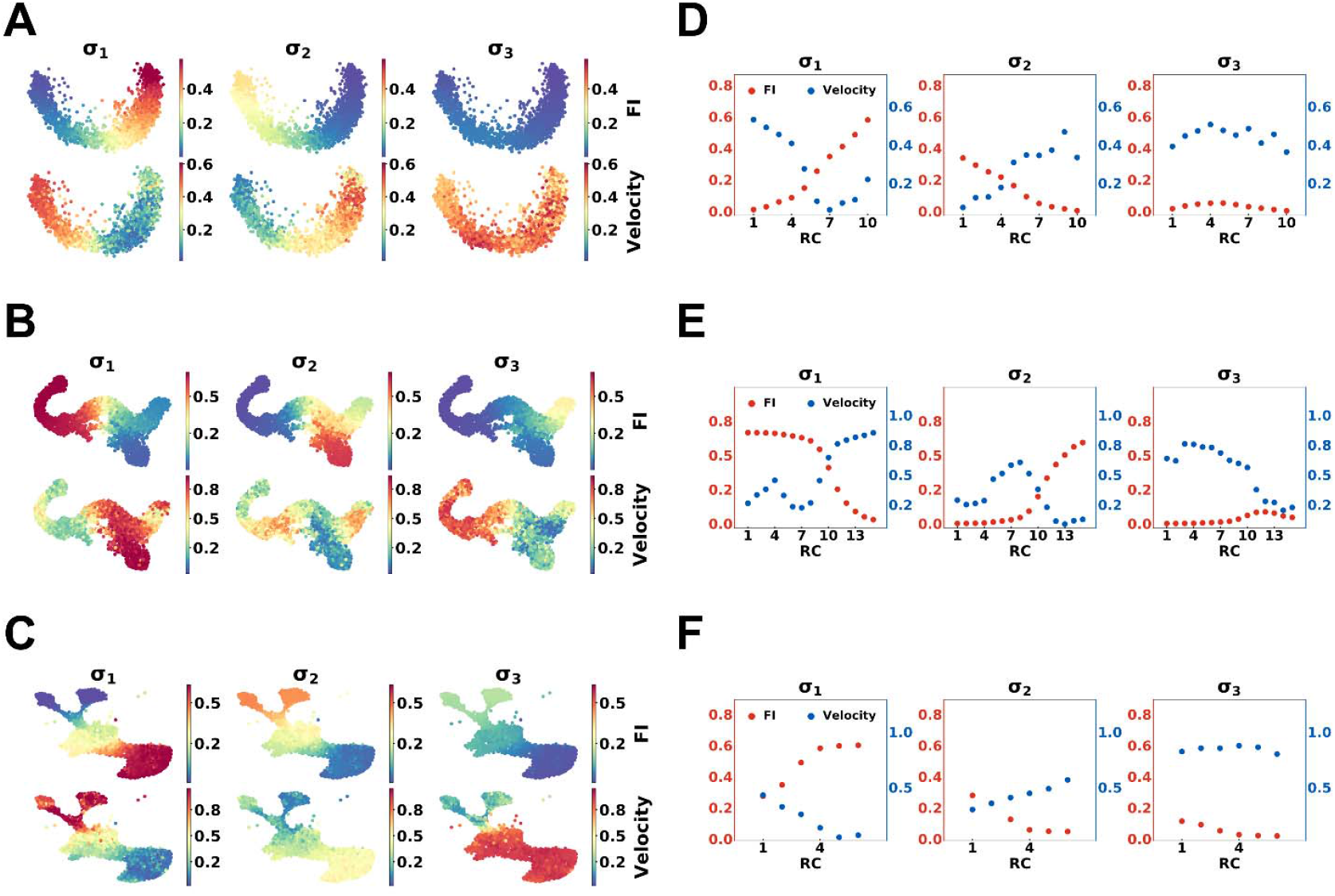
Relationship between the parameter velocity and the Fisher information reveals the action constraint on transition paths. (**A**) Cell-wise distributions of the Fisher information and the parameter velocity in the dentate gyrus neurogenesis dataset. Each dot represents a single cell, with color indicating the value of the Fisher information (top) or parameter velocity (bottom). For each cell, the Fisher information and parameter velocity of each parameter are sum-normalized across all parameters. (**B**) Same as that in panel (A) except for the pancreatic endocrinogenesis dataset. (**C**) Same as that in panel (A) except for the zebrafish pigmentation dataset. (**D**) Relationship between the Fisher information and the parameter velocity along the transition path in the dentate gyrus neurogenesis dataset. The Fisher information of a given parameter is computed by averaging over neighboring cells of each RC point, and the parameter velocity is computed in the same manner. The Fisher information and parameter velocity are sum-normalized across parameters. (**E**) Same as that in panel (D) except for glucagon-producing α-cell branch of the pancreatic endocrinogenesis dataset. (**F**) Same as that in panel (D) except for the melanophore branch of zebrafish pigmentation dataset.

We further calculated the average Fisher information and parameter velocity of neighboring cells for each RC along the transition paths (Fig. 6D-F). Two major characteristics emerge from this analysis. First, parameter velocity is low when Fisher information is high, indicating that stiff parameters do not exhibit rapid variation during CST. This is consistent with our previous result showing that stiff parameters have lower fluctuations (Fig. 5B). Second, parameter velocity can fluctuate when Fisher information is low. For example, in the pancreatic endocrinogenesis dataset, while Fisher information of σ_2_ remains low across all RC points, its parameter velocity fluctuates, being high in the first half of the transition path and low in the second half (Fig. 6E). In another word, the velocity of the sloppy parameter is less restricted compared to that of the stiff parameter.

We can interpret these results from the perspective of action on the Riemann manifold. Consider the action along the transition path

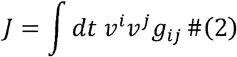

which is the integral of the square of information velocity along the path. Information velocity 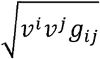 represents the variation rate of p istribution, i.e., cell state. This action has a lower bound related to the length of the path *L*

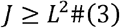

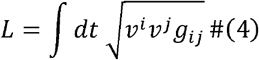

The paths which minimize this action to the lower bound are the geodesics on the manifold that satisfies

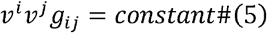

(*25*). From the definition, we can see that to minimize the action, which is equivalent to keeping the information velocity constant along the path, the velocity and metric must vary in a complementary way: *g*_*ij*_ must be small where *v*^*i*^*v*^*j*^ is large, and vice versa. This imposes a constraint on the parameter velocity in turn. For sloppy parameters whose influence on information velocity is minimal, their parameter velocities are less restricted. However, for stiff parameters, the parameter velocity is constrained to low levels because variations in stiff parameters can significantly alter the cell state. Therefore, we can conclude that cells tend to be “sloppy,” meaning they tend to minimize information velocity. When a parameter’s Fisher information changes, its parameter velocity typically varies in the opposite direction.

In CST processes, the situation is complex. As we have discovered that the information velocity transiently increases at the transitory stage in CST from one cell state to another state (*20*). The increase of information velocity reflects the rapid variation of cell state. In CSTs involving branching into different lineages, the trend in information velocity is more complicated, as it involves the regulation of different stiff parameters or interactions between various GRN modules. Nevertheless, these findings suggest that the CST process generally progresses in a way that seeks to minimize action most of the time, possibly to conserve energy. However, around the transitory stage, this principle is not upheld, as the cell state undergoes significant changes and becomes highly sensitive to external stimuli, consuming substantial energy. From a control perspective, cells may prefer to adjust sloppy parameters to avoid state transitions unless such transitions are unavoidable. Results for the other parameters and branches are shown in Fig. S17 and S18. The robustness tests of the Gaussian embedding hyper-parameters are shown in Fig. S19-27.

## Discussion

Bifurcation diagram is commonly used to illustrate CSTs. However, analyzing the bifurcation of high-dimensional systems is challenging, as it typically requires a clear definition of cell dynamics. The variation of FIM offers a new perspective for analyzing complex state transitions. Other than analyzing CST in the dynamics space, we mapped the cell dynamics in CST to parameter space (Fig. 7) and characterized CST processes with variation of the FIM spectrum, particularly the switch of stiff parameters. The augmentation of stiff parameters in some branching processes may correspond to an increase in dynamic modes within the behavioral space.

**Fig. 7.**
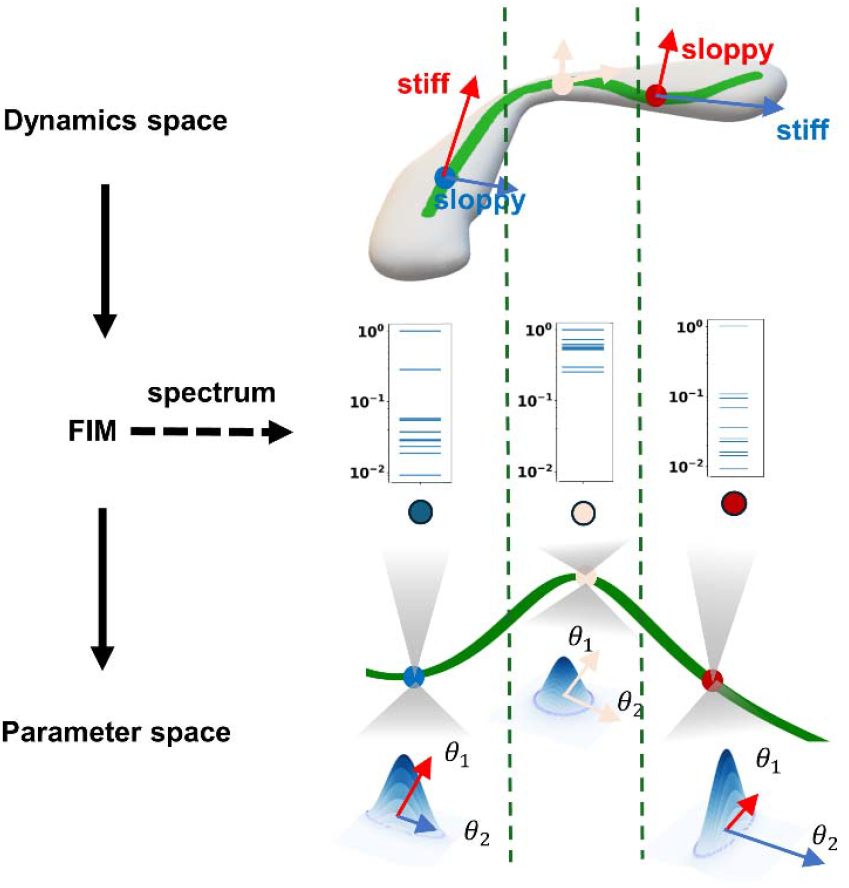
Analyzing high-dimensional dynamics of CST (bottom) through mapping onto parameter space (top) using the FIM. The two green curves in two spaces represent the same trajectory, with points of matching colors denote the same model state. The transition can be studied by analyzing the FIM eigenvalue spectrum. Along the path, the stiff direction switch between (red) and (blue). During this switching, the second eigenvalue of FIM becomes comparable to the first one, illustrating terminal states with a dominant direction and transitory state where multiple directions become similarly informative. As a schematic example, each point corresponds to a two-dimensional Gaussian distribution in the parameter space, with arrow length at each point indicating the magnitude of the Fisher information in that direction.

Given the limitations of defining paths directly from experimental data, we adopted an alternative approach to analyze action constraints. We examined the velocities of stiff and sloppy parameters, interpreting our observations from both control and action perspectives. Compared to that in classical mechanics, this “approximate least action principle” fails to hold around the transitory stage. However, this failure coincidentally uncovers a common characteristic of CST: stiff parameters exhibit increased fluctuations and velocities at the transitory stage. Establishing a more general principle will require both the advancement of techniques to capture real trajectories experimentally and the development of theoretical tools from information geometry.

It is important to note that Fisher information is not directly related to causal relationships; however, we can still gain insights into system control by mapping it onto parameter space. In our analyses, we used linear decomposition of the FIM to study the variation of stiff directions. Nonetheless, stiff directions can be certain nonlinear combinations of individual parameters. Further development of nonlinear methods for analyzing sloppiness from the FIM could enhance our understanding (*52*).

In CST, GRNs are likely dynamic, with variations in regulation strength guiding critical transitions (*38, 39*). Even though the collective coordinates like Eigengene and Hotspot modules can reflect the organization of GRN, they cannot fully capture the direct interactions between gene modules or the regulatory strength among genes. And labeling a gene as “stiff” indicates that it has a significant impact on biological processes and plays a dominant role. Such genes are more likely to interact with others, leading to larger non-diagonal elements in the FIM corresponding to those gene pairs. Nonetheless, this is not a formal definition of Fisher information concerning gene interaction. A potential approach would involve inferring GRN and calculating the Fisher information of regulatory strength between genes.

## Methods

### Dataset and preprocessing

The scRNA-seq dataset of zebrafish pigmentation is available on GEO via accession GSE131136 (*41*). We filtered out genes expressed in less than 10 cells and selected 600 highly variable genes for analysis. For simplicity, we focused on specific transition processes and filtered out other cell types not involved in these processes for calculation considerations. The focused differentiation processes are proliferating progenitors to melanophore, xanthophore, and Schwann cell. Cells annotated as other cell types were removed. The dentate gyrus neurogenesis is obtained from the GEO website with accession number GSE95753 (*42*). We selected 678 genes that display switch-like behaviors by using the approach described in (30). In this dataset, we first analyzed the single-lineage CST process from radial glia cell to mature granule cell for comparison with multiple-lineage CST processes like zebrafish pigmentation. The pancreatic endocrinogenesis dataset is obtained from GSE132188 (*48*). We selected 470 genes that display switch-like behaviors by using the approach described in (30). In this dataset, we include both branches of glucagon-producing α-cells and β-cells.

### Single cell information manifold (SCIM)

SCIM is a method we previously developed to quantify single cell manifolds (*20*). Briefly, SCIM employs a machine learning method for graph embedding (40,41), called Gaussian embedding, to transform each cell’s *N*-dimensional scRNA-seq data into an *L*-dimensional (*L* < *N*) embedding represented by the parameters (means and variances) of a multivariate diagonal Gaussian distribution. To do so, we first constructed a k-nearest neighbor cell graph (*k*-NN) from data. Secondly, a neural network is trained on the sampled triplets which contain the neighboring information of nodes (cells) in the cell graph by minimizing an energy-based ranking loss (ℒ)

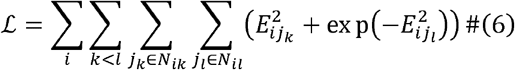

where *N*_*ik*_ represents the set of all nodes *j*_*k*_ satisfying the condition that the length of shortest paths on the cell graph between node *i* and *j*_*k*_ equal *k* and the energy *E*_*ij*_ represents the Fisher distance between nodes *i* and *j*

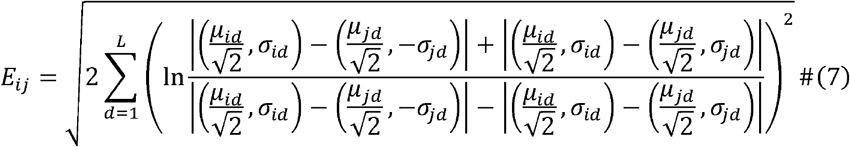

When minimizing the energy-based loss, the Fisher distance between nodes that are close in the gene expression space decreases, whereas the Fisher distance between nodes that are farther apart tends to increase. Intuitively and empirically, SCIM preserves the local and global geometric structures of the data manifold by maintaining the relative distances between node pairs (10).

In this study, the neural network has two hidden layers (256-128 nodes) equipped with ReLU activation, a linear μ-head, and a linear σ-head equipped with ELU activation. To ensure positive variance, we added a unit offset to the σ-head. All neural networks used for Gaussian embedding were trained for 400 epochs using Adam with a learning rate of 0.001. Gaussian embedding involves three main hyperparameters: the number of nearest neighbors *k*_*nei*_ used to construct the *k*-NN graph, the maximum hop number *K* in the neighborhood graph considered in the loss function, and the latent embedding dimension *L*. In the robustness tests of model hyperparameters in this paper, we performed tests across *k*_*nei*_=10,20,30, 2,3, and *L*=3,6,10,15.

### Eigengene

To investigate the potential gene regulatory mechanisms and characteristics underlying CST, we employed the Eigengene method introduced by Langfelder in 2007 (*45*). Based on correlations between genes, we calculated the gene co-expression patterns. Using gene–gene correlation as a distance measure, we performed hierarchical clustering to partition genes into different modules. The Eigengene of each module is defined as the first principal component (PC1) of that module, serving as the representative expression profile of the gene module.

### Hotspot

Another method for identifying gene modules that capture shared gene-expression features is Hotspot (*46*). This algorithm computes a local expression magnitude of each gene and a neighborhood-based metric to decide whether the gene expression pattern is informative i.e., if it is associated with the *k*-NN structure. It also calculates the similarity between genes. For example, by constructing a *k*-NN graph based on distances among cells in latent space, physical space, phylogeny, etc., one can determine whether a gene expression pattern is related to the cell’s position in latent space, physical location, or phylogeny.

Concretely, given a weighted *k*-NN graph, for each gene *x*, the autocorrelation on the graph is defined as

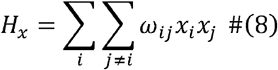

where *ω*_*ij*_ denotes the weight of the edge connecting cells *i* and *j* in the graph, and *x*_*i*_ denotes the standardized expression of gene *x* in cell *i* (zero mean, unit variance). If a gene *x* has a relatively high autocorrelation on the graph — indicating that its expression is closely tied to the graph structure—it is considered informative.

Hotspot computes pairwise association scores for all informative genes as follows

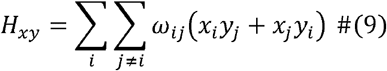

This association score describes how the expressions of genes *x* and *y* are influenced by the graph structure. Unlike the classic Pearson correlation, this association score is not the inner product of two gene profiles; rather, it measures how the expression of gene *x* in a cell depends on the expression of gene *y* in neighboring cells, and vice versa. Gene pairs with larger absolute association values represent stronger similarity in expression patterns. By treating this similarity as a distance measure, hierarchical clustering is performed on all informative genes to partition genes into different modules. The PC1 value of a module is referred as the module’s Hotspot coordinate and serves as the representative expression of that gene module.

### Eigenvalue spectrum of FIM

For probability distribution with parameter *θ*, its FIM is defined as

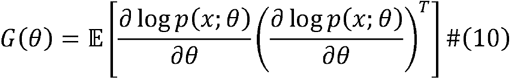

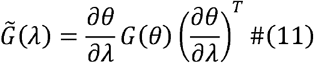

where *p*(*x*; *θ*) is probability density function and 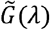 is the FIM of another parameter λ computed through coordinate transformation from parameter *θ* to parameter *λ*. This allows us to observe the spectrum of the FIM’s eigenvalues in different spaces.

In (μ,σ) space, the FIM is diagonal, so the eigenvalues are simply the diagonal entries. We trained a neural network to learn the coordinate transformation using the auto-gradient of the neural network. In principle component analysis (PCA), Eigengene, Hotspot, or other coordinate systems, we applied symmetric eigen decomposition to compute eigenvalues and eigenvectors. Since FIM is regarded as a metric matrix, the scale of its eigenvalues indicates how rapid the distribution changes.

### Transition path analysis

Although single-cell data typically consist of snapshot data lacking dynamic information, the transition path can still be inferred based on prior knowledge and reasonable assumptions. To better illustrate the variation of Fisher information during the transition process, we simulated transition trajectories on each scRNA-seq data. The simulation was conducted using a transition kernel defined on the *k*-NN graph. We employed an EM-like algorithm to derive a smooth average transition path (*40*). We used the cell-cell transition matrix provided by scVelo package as the transition kernel (*53*)

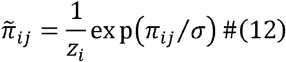

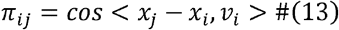

Here, *x*_*i*_ represents cell *i, v*_*i*_ represents the RNA velocity of cell *i*. The cosine similarity *π*_*ij*_ between potential cell state and RNA velocity was computed. The transition probability 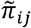 was then determined by applying an exponential kernel.

When simulating trajectories, we first utilized PAGA or cell type annotations to divide cells into several clusters, specifying an initial cluster and a final cluster. We then calculated the average cell of the starting cluster and selected the 200 cells nearest to it as starting points to generate 200 trajectories. Each trajectory that ends in the final cluster (terminated after remaining in the final cluster for 10 steps) is used for averaging if its length is less than 200. For averaging, we employed an iterative EM-style hard-assignment procedure. In each iteration, trajectory time points were assigned to the nearest spline samples, and the spline control points were updated through weighted averaging and re-fitting, continuing until the fitting error converged.

Given a transition path discretized by RCs, we separated the cells into Voronoi grids by identifying the nearest RC for each cell. The Fisher information of the *i-th* RC was defined as the mean Fisher information of cells within the *i-th* grid. The projected velocity along path was also defined in this way. The standard deviation and coefficient of variation of projected velocity along the path was defined as the std and CV of projected velocity in each cluster. Similarly, the standard deviation (std) and coefficient of variation (CV) of parameter velocity along the path was defined in this manner.

For spectrum analysis, we selected 10 RCs for dentate gyrus neurogenesis; 15 RCs for both branches of glucagon-producing α-cells and β-cells in endocrinogenesis; 10 RCs for the zebrafish melanophore-branch; and 8 RCs for the zebrafish xanthophore and Schwann cell branches.

### Gene Fisher information analyses

Given Fisher information metrics in (*μ,σ*) space, we pulled it back to gene expression space

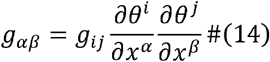

where *x*^*α*^ denotes the *α*-element of the gene expression vector, *θ*^*i*^ denotes the *i*-element of the latent state vector in (*μ,σ*) space. The diagonal element *g*_*αα*_ is defined as the gene Fisher information of the gene α.

We trained a multilayer perceptron (MLP) with two hidden layers (128-64, ReLU activations) to map from gene expression space to (*μ,σ*) space. The Jacobian was obtained by automatic differentiation of the MLP. These MLPs were trained using the Adam optimizer with a learning rate of 0.001.

We analyzed the variation of Fisher information distribution of individual genes within each cell cluster or cell type. We selected the top 20 genes with the highest average Fisher information across all cells for dot plot visualization (Fig. 3A). For each selected gene, we calculated its Fisher information in all cells and normalized it by its maximum Fisher information across all cells to obtain a relative value. The relative Fisher information distribution of individual gene was fitted using kernel density estimation (KDE).

We also examined the distribution of the average Fisher information for all genes (on a log scale) at each RC. The log Fisher information of each gene at a given RC was defined as the mean log Fisher information of that gene across the 50 nearest cells to that RC. We fitted this distribution with a two-component Gaussian Mixture Model and used the t-test distance to measure the separation between the two Gaussian components, while the Gini coefficient was employed to assess the dispersion of the distribution. Denoting the 2 components as 𝒩 (*μ*_1_,*σ*_1_^2^) and 𝒩 (*μ*_2_,*σ*_2_^2^) the separation and Gini coefficient are defined as follows

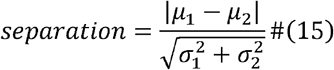

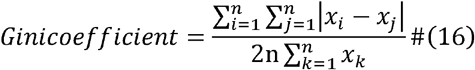

where *x*_*i*_ are *i*.*i*.*d* samples from a distribution supported on positive real numbers. Note that the distributions of log Fisher information are supported on negative real numbers; thus, we multiplied the data by -1 before calculating the Gini coefficients.

### Clustering and visualization of stiff direction

Before clustering, we addressed the sign’s ambiguity issue, as the reverse direction of a stiff direction is still considered a stiff direction. This can affect the calculated distances between vectors, thereby influencing the hierarchical clustering results. The optimal sign for each eigenvector is determined by solving the following optimization problem

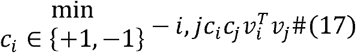

where *v*_*i*_ denotes the eigenvectors. Since the vectors are unit vectors, we minimize their negative correlation in the optimization process; this serves as an indicator for determining the direction of each vector. This formulation constitutes a binary quadratic optimization (BQO) problem. We employ the Gurobi Optimizer to obtain the solution, utilizing the commonly used branch-and-bound algorithm from integer programming (*50, 54*).

Hierarchical clustering is a class of distance-based clustering methods (*55*), with bottom-up agglomerative hierarchical clustering being the most prevalent variant. We adopt an adaptive threshold selection rule based on a probabilistic perspective. As the dimensionality of the vector space increases, the angle (cosine distance) between two randomly selected vectors in the N-dimensional space is more likely to approach 90 degrees. Consequently, we use the critical value corresponding to the 95% confidence level of the cosine distance distribution between two random unit vectors as the threshold. If the cosine distance between two vectors falls below this threshold, we consider there to be sufficient confidence to group them together. The probability of this event is proportional to the area of the corresponding region on the N-dimensional unit sphere, and the formula for calculating this probability is provided below

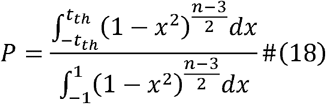

where *x* represents the cosine of the angle *t*_*th*_ represents the cosine of the angle threshold. Wesolve *t*_*th*_ by set *P*= 0.95 and get the distance threshold *D*_*th*_ = 1− *t*_*th*_.

To project these directions onto a specific 2D visualization space (e.g., UMAP), we trained a MLP, denoted as *ϕ*, to map coordinates from the parameter space to the target space. Each arrow in Fig 4A is derived from the finite difference calculated by perturbing a point *x*_*i*_ in parameter coordinate along its corresponding stiff direction *d*_*i*_ by a small step size *δ* and then mapping the result using the MLP

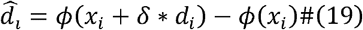

The orientation of the plotted arrow corresponds to the direction of 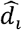. The length of the arrow is proportional to the size of the stiff direction cluster.

### Fluctuation and velocity of parameters

We used the squared coefficient of variation *CV*^2^ of parameters at each RC point to access the fluctuations of stiff and sloppy parameters following the same idea used for computing *projection change* in (*35*).

The parameters considered here represent the coordinates defined by the eigenvectors of the FIM in PCA space, obtained as follows

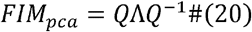

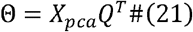

where, *Q* is an orthogonal matrix, whose columns are the eigenvectors of *FIM*_*pca*_,Λis a diagonal matrix containing the corresponding eigenvalues (Λis also the FIM of the parameters *λ*), and *X_pca* is the principal component score matrix of cells obtained by projecting the highly variable gene expression matrix onto the leading principal components (top 50). *FIM*_*pca*_ refers to the FIM in PCA space. Here, 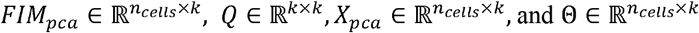, where *ncells* is the number of cells in the dataset and *k* is the number of principal components considered (*k* = 50). Each element of Θ represents a cell’s new parameter, obtained as the projection of its PCA coordinates onto a Fisher eigenvector.

For each RC point, we selected the 30 nearest neighboring cells as samples to calcu,ate *CV*^2^, based on the assumption that cells located close to an RC point share the same cell state and are governed by a similar set of parameters. As an example, the *CV*^2^ of the stiffest parameter at the first RC point *RC*_1_is computed as follows

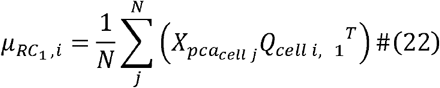

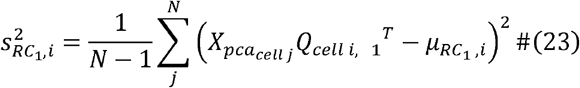

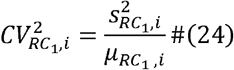

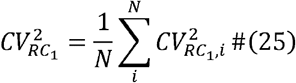

where, *Q*_*cell i*,1_ denotes the eigenvector corresponding to the largest eigenvalue of *FIM*_*pca*_ for *celli*, 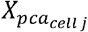 denotes the PCA coordinates of neighboring *cell j*, and *N* is the number of neighboring cells, here *N* = 30.

For Gaussian embedding, the parameter velocity of (μ,σ) can be computed with RNA velocity

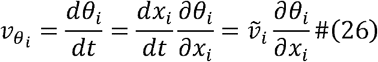

where, 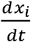 is the original RNA velocity for gene *i* and 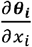 is calculated using Pytorch autograd function numerically from Gaussian embedding model.

Each individual cell is assigned a set of parameters. For each parameter, such as ***θ***_***i***_, the sum-normalized Fisher information and its velocity are calculated as follows

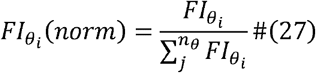

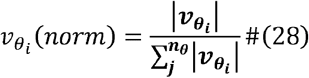

Here, *n*_*θ*_ is the number of parameters for each cell, and 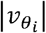 denotes the absolute value of the velocity of the parameter *θ*_*i*_.

## Acknowledgement

W.W. is supported by National Natural Science Foundation of China Grants No.12247104. L.Z. was supported by the National Key Research and Development Program of China 2024 YFA0919500 and National Natural Science Foundation of China (No. 12225102, T2321001, and 12288101). All data needed to evaluate the conclusions in the paper are present in the paper and the Supplementary Materials. The codes are available at https://github.com/wwklab/sloppy-SC.

## Author contributions

Conceptualization: WW; Methodology: JY, YW, HX, MH; Investigation: JY, YW, HX; Visualization: JY, YW, HX; Supervision: LZ, WW; Writing—original draft: JY, YW, HX, MH, LZ, WW; Writing—review & editing: LZ, WW

